# Transcriptomic responses of Mediterranean sponges upon encounter with seawater or symbiont microbial consortia

**DOI:** 10.1101/2023.11.02.563995

**Authors:** A. Marulanda-Gomez, M. Ribes, S. Franzenburg, U. Hentschel, L. Pita

**Affiliations:** RU Marine Symbioses, RD3 Marine Ecology, GEOMAR Helmholtz Centre for Ocean Research Kiel, Germany; Institut de Ciències del Mar, ICM – CSIC, Spain; Research Group Genetics and Bioinformatics/Systems Immunology, Institute of Clinical Molecular Biology, Christian-Albrechts-Universität Kiel, Germany; Christian-Albrechts-Universität Kiel, Germany

**Keywords:** animal-microbe interactions, microbial consortia, HMA-LMA sponges, immune receptors, RNA-Seq, differential gene expression, symbiosis

## Abstract

Sponges (phylum Porifera) constantly interact with microbes from the water column while filter-feeding and with the symbiotic partners they harbor within their mesohyl. Despite early observations on differential uptake between symbiont and seawater bacteria, it is still poorly understood how sponges discriminate between different microbial consortia. Initial evidence of the diverse repertoire of sponge immune receptors suggests their involvement in specific microbial recognition, yet experimental data is still scarce. We characterized the transcriptomic response of two sponge species, *Aplysina aerophoba* and *Dysidea avara*, upon incubation with two different microbial consortia, which were either enriched from ambient seawater or extracted from *A. aerophoba.* The sponges were sampled after 1 h, 3 h, and 5 h for RNA-Seq differential gene expression analysis. *D. avara* showed higher expression levels of genes related to immunity, ubiquitination, and signaling when incubated with *A. aerophoba* symbionts, than in incubations with seawater microbial consortia. Interestingly, the different bacteria consortia triggered changes in Nucleotide Oligomerization Domain (NOD)-Like Receptors (NLRs) gene expression in *D. avara*. We here provide the first experimental evidence for NLRs playing a role in distinguishing between different microbes in a sponge. In contrast, the response of *A. aerophoba* involved comparatively few genes and lacked genes encoding for immune receptors. This indicates that *A. aerophoba* is less responsive to microbial encounters than *D. avara*. Our study further reveals different transcriptomic responses between the two sponge species to microbes. The study of sponge responses to microbes aids in understanding the evolution of immune specificity and animal-microbe interactions.

**Significant statement:** Animals rely on components of the immune system to recognize specific microbes, whether they are pathogens, food, or beneficial symbionts. However, in marine invertebrates, the mechanisms of microbial discrimination and specificity are not well understood. Our work suggests that:(i) the transcriptomic response by the sponge can be scaled according to the type of exposure, (ii) the response to microbial encounters is species-specific and (iii) NLRs seem to have a prominent role in the differential response to microorganisms, whether symbionts or food bacteria.

## Introduction

Over the last decades, animals were recognized as “metaorganisms” or “holobionts” which encompass the multicellular host and its microbial symbionts, such as bacteria, archaea, viruses, fungi, and algae, (Bosch & McFall-Ngai, 2021; González-Pech et al., 2023; Rosenberg & Zilber-Rosenberg, 2018; Stévenne et al., 2021). Symbionts participate in the general fitness of the host by contributing to developmental cues, nutrient provision, potential metabolic expansion, and defensive traits (Carrier & Bosch, 2022; Gilbert et al., 2015; McFall-Ngai et al., 2013; Wein et al., 2019). Microbes thus provide adaptive advantages and shape animal evolution (Rosenberg & Zilber-Rosenberg, 2016; Roughgarden et al., 2018). These close and complex host-microbe interactions required fine-tuned communication between partners which is now known to be orchestrated by the host immune system (Berg et al., 2019; Dierking & Pita, 2020; Ganesan et al., 2022; Horak et al., 2020).

The immune system has a dual function of defending the animal against potentially harmful intruders and at the same time, establishing and maintaining interactions between the host and its symbiotic microbes (Eberl, 2010; Gerardo et al., 2020). How does immunity differentiate pathogens to be eliminated from symbionts to be acquired/maintained, and how does it safeguard guard homeostasis and equilibrium within the host? In both contexts, animals sense microbe-associated molecular patterns (MAMPs), such as lipopolysaccharide, peptidoglycan, or flagellin, via pattern recognition receptors (PRRs) (Janeway & Medzhitov, 2002). However, the encounter to a pathogenic microbe elicits inflammatory responses to eliminate the intruder, whereas the interaction with symbionts results in tolerance and colonization (Chu & Mazmanian, 2013; Gerardo et al., 2020). Thus, the immune system is able to specifically distinguish between symbiotic, non-symbiotic and pathogenic signals.

Many invertebrate groups such as hydrozoans, cnidarians, mollusks, and echinoderms present large and strikingly complex repertoires of PRRs (Buckley & Rast, 2015; Hamada et al., 2013; Lange et al., 2011; Neubauer et al., 2016; Zhang et al., 2011). The diversity of these receptors contributes to microbial detection by the host, and potentially plays a role in microbial discrimination (Jacobovitz et al., 2021; Neubauer et al., 2016; Saco et al., 2020; Seneca et al., 2020). For example, the coral *Montipora aequituberculata* responds to potentially pathogenic and commensal bacteria *Vibrio coralliilyticus* and *Oceanospirillales* sp., respectively by regulating the expression of Toll-like receptors and via differential upregulation of G protein– coupled receptors (van de Water et al., 2018). On the other hand, the freshwater snail *Biomphalaria glabrata* recognizes different pathogens by different sets of PRRs belonging to the calcium-dependent lectin family and via enzymes and non-canonical immune components, like extracellular actin (Tetreau et al., 2017).

As arguably the earliest branching metazoans (Redmond & McLysaght, 2021; Turner, 2021), sponges (phylum Porifera) offer the opportunity to study the evolution of immune specificity and animal-microbe interactions. As active filter-feeders pumping thousands of liters of seawater per day through their aquiferous system, sponges constantly encounter microbes from the seawater, but, at the same time, harbor specific and complex microbial communities within their mesohyl matrix (Thomas et al., 2016; Webster & Thomas, 2016). Based on the density and diversity of their symbionts, sponges are classified as high microbial or low microbial abundance (HMA and LMA) species. HMA sponges contain three to four order of magnitude more microbes than LMA sponges (Bayer et al., 2014; Hentschel et al., 2006; Moitinho-Silva et al., 2017; Pankey et al., 2022). This long-recognized dichotomy in sponge-microbe symbiosis reflects particular signatures in the structure and persistence of the symbiosis as well as physiological differences such as density of the mesohyl and pumping rates (Gloeckner et al., 2014; Maldonado et al., 2012; Morganti et al., 2021; Weisz et al., 2008)). Additionally, LMA sponges are enriched in genes involved in microbial sensing and in host defense, such as SRCRs, NLRs, nucleosome-binding proteins, and bactericidal permeability-increasing proteins compared to HMA sponges (Germer et al., 2017; Ryu et al., 2016). The expression of immune-related genes upon different stimuli thus depends on the microbial abundance and diversity associated with the sponge, but can also be species-specific (Campana et al., 2022; Pita et al., 2018; Posadas et al., 2021). These observations suggest that the HMA-LMA status, as well as specific species traits, may affect how the sponge immune system responds to microbial cues.

Early experimental evidence showed that sponges preferentially take up seawater microbial consortia (i.e., bacterioplankton) over sponge symbiont consortia (Wehrl, 2006; Wehrl et al., 2007; Wilkinson et al., 1984). This was taken as evidence that the animal can differentiate between different microorganisms. This differentiation may derive from both host and microbial features. Recently, sponge symbionts were shown to evade phagocytosis by the expression of eukaryotic-like proteins containing ankyrin repeats which silence conserved components of phagocytosis and immune signaling (Jahn et al., 2019). On the host side, the high diversification of PRRs in sponges (Hentschel et al., 2012; Riesgo et al., 2014; Srivastava et al., 2010; Yuen et al., 2014) suggest their potential to recognize different and specific microbial ligands (Degnan, 2015). Experimental evidence supports the activation of PRR gene expression upon encounter with microbial elicitors (e.g., Wiens et al. 2005; Yuen 2016; Pita et al. 2018; Schmittmann et al. 2021), but it remains to be shown if (and how) the expression patterns of PRRs may be involved in specific immune responses to different microbes.

Our study aims to better understand the underlying molecular mechanisms of bacterial discrimination in sponges. We characterized the host response upon encounter to seawater- and sponge-derived microbial consortia by RNA-Seq differential gene expression analysis. Specimens of *Aplysina aerophoba* (HMA) and *Dysidea avara* (LMA) were incubated with either microbial consortia enriched from natural seawater or with a sponge-associated symbiotic consortia. Sponge symbiont consortium was obtained from *A. aerophoba* by differential centrifugation, a physical separation used for enrich sponge symbiotic fractions because sponge symbionts remain unculturable (Schmittmann et al., 2022; Markus Wehrl et al., 2007). We collected samples at 1h, 3h, and 5h from the start of the incubation. We hypothesized that (i) both sponges will rely on differential expression of PRRs for microbial discrimination, (ii) differentially-expressed genes will show lower expression levels upon symbiotic (“self”) than seawater microbial consortia (“non-self”) treatment and (iii) that the HMA-LMA status of the host sponge will influence the different transcriptomic response between microbial encounters.

## Material and Methods

### Sponge collection

Specimens of the Mediterranean sponge species *Aplysina aerophoba* (Nardo, 1833) and *Dysidea avara* (Schmidt, 1862) were collected via SCUBA diving at the coast of Girona (Spain) in March 2015 (42.29408 N, 3.28944 E and 42.1145863 N, 3.168486 E; respectively). Sponges were then transported to the Experimental Aquaria Zone (ZAE) located at the Institute of Marine Science (ICM-CSIC) in Barcelona (Spain) and were placed in separated 6 L aquaria in a flow-through system with direct intake of seawater. Temperature and light conditions were set up mimicking natural conditions. Sponges were maintained under these conditions during 10-12 days for acclimation.

### Experimental setup

The experiment was conducted consecutively for each sponge species (end of March for *A. aerophoba*, beginning of April for *D. avara*). Before the microbial exposure experiments, sponges were kept overnight in 1 µm-filtered seawater and an additional 0.1 µm-filter was applied for 3 h before the experiments with the aim to reduce microbial load in seawater to a minimum. The flow-through was stopped during the experiment, but small aquarium pumps (Eheim Gmbh & Co.) ensured the mixing of the water in the aquarium. Sponges were incubated with either microbial seawater consortia or symbiont consortia that had been prepared following the protocols below. The concentration of these stock microbial consortia was estimated via flow cytometry (see details in supplementary information, Text S1 and Fig. S1) and adjusted to reach 10^5-6^ bacteria mL^-1^ final concentration in the experimental tanks. Sponge specimens that were actively pumping, as visually assessed by the presence of an open oscula, were randomly assigned to each treatment (n = 5 individuals per treatment). For each individual, tissue samples were collected at 1 h, 3 h, and 5 h after adding the microbial consortia to the experimental tanks, then placed in RNAlater at 4°C overnight, and stored at -80°C until processing.

### Symbiont consortia preparation

The *A. aerophoba*-symbiont fraction was obtained as described in Wehrl et al. (2006). Briefly, 20 g of sponge tissue from living individuals that had been cleaned off debris was rinsed in sterile, ice-cold Ca- and Mg-free artificial seawater (CMFASW) with EDTA (as in (Rottmann et al., 1987)), incubated for 30 min at 4°C, and then homogenized with a mortar and pestle. After filtration through 100 µm-Nitex, the suspension was centrifuged twice at 4°C, 400 g for 20 min to remove sponge cells, which remained in the pellet. The supernatants were combined and centrifuged at 4°C, 4000 g for 20 min to obtain a bacterial pellet. This pellet was washed twice in ice-cold CMFASW and recovered again by centrifugation. Finally, the bacterial pellet was resuspended in sterile ice-cold CMFASW. Symbiont extraction from *D. avara* was not possible because this species represents an LMA sponge and we could not obtain enough microbial extracts for the incubations.

### Seawater microbial consortia preparation

Seawater microbial consortia were enriched from seawater from the aquaria setup (a flow-through system with direct intake of natural seawater), following the protocol by Wehrl et al. (2006). In short, Marine Broth 2216 media was added to 10 L of seawater to a final concentration of 15 mg L^-1^. The enriched seawater was incubated in the dark overnight at ambient temperature and gentle shaking. Aliquots of the enriched seawater were then sampled, and bacteria were recovered by differential centrifugation (4°C, 4000 g for 20 min), then washed twice, and re-suspended in sterile, ice-cold CMFASW.

### Sponge RNA extraction, sequencing, and *de novo* transcriptome assembly

Total RNA from 30 samples were extracted for each species following the methods in Pita et al. (2018), but only 22 samples of *D. avara* pased the quality checks (i.e., RIN > 8 in Experion, Bio-Rad, USA). In short, 500 ng of total RNA were used for library construction with the TruSeq stranded mRNA library prep kit (Illumina, Inc., USA), including a poly-A enrichment step. Paired-end sequencing (150 bp) was performed on a NovaSeq S2 system (Illumina, Inc., USA) at the Competence Centre for Genomic Analysis (CCGA; Kiel, Germany). Raw paired-end reads were trimmed and filtered to remove adapters and low-quality reads in Trimmomatic-v0.39 (Bolger et al. 2014). Prokaryotic and microbial eukaryotic reads were filtered in the classifier kaiju-v1.6.2 (Menzel & Krogh, 2015). All samples were used to construct a *de novo* assembly per each sponge species in Trinity-v2.10.0 (Haas et al., 2013). Quality check and completeness of the assemblies were assessed by statistics performed in TransRate-v1.0.2 (Smith-Unna et al., 2016), and by comparing the assemblies against the metazoan-reference data in BUSCO-v3 (Simão et al., 2015).

### Annotation, gene quantification, and differential gene expression analysis

Functional transcriptome annotation was performed following Trinotate-v3.2.0 (Haas et al., 2013). Contigs with blastx or blastp matches to Bacteria, Archaea, or Virus, as well as those annotated as ribosomal RNA were removed from the *de novo* assembly. Gene (i.e., trinity components) abundance was estimated based on RSEM bowtie2 quantification-v1.3.3 (Langmead et al., 2009; Li & Dewey, 2014). Differential gene expression analysis was performed separately for each time point (i.e., 1 h, 3 h, and 5 h) in edgeR (Robinson et al., 2009) as implemented in Trinity-v2.10.0 (Haas et al., 2013) with default parameters. Differentially expressed genes (DEGs) in pairwise-treatment comparisons were defined by False Discovery Rate-corrected (FDR) p-value < 0.005 and log2|change| ≥ 2 (i.e., four-fold change) as in (Pita et al., 2018; Wu et al., 2022).

## Results

We characterized the transcriptomic response of the Mediterranean sponges *A. aerophoba* and *D. avara* to either seawater microbial consortia or to symbiont consortia extracted from *A. aerophoba* tissue. We followed upon the initial work by Wehrl et al. (2007), who observed lower uptake rates of symbionts than seawater bacteria in *A. aerophoba*.

### Reference transcriptome assembly

We sequenced 29 samples of *A. aerophoba* and 22 samples of *D. avara* corresponding to 3-5 biological replicates per treatment within 1h, 3h, and 5h (Table 1). The number of paired-end Illumina reads generated in this study is summarized in Supplementary Table S1. BUSCO assessments revealed that the *de novo* reference transcriptomic assembly of *A. aerophoba* generated in this study contained 71.4 % of the 902 BUSCO Metazoan core genes, with 76.6 % of the genes found as complete. The reference transcriptome assembly for *D. avara* consisted of 78.2% of the BUSCO Metazoan core genes, with 82.9% of these genes found as complete, suggesting this reference transcriptome is more complete than the reference in Pita et al. (2018). All statistics of the reference assemblies generated in this study are summarized in Supplementary Table S2. Overall, 68.89 ± 0.21% and 84.37± 17% (average ± standard error) of the reads in each sample aligned to the *de novo*-assembled reference transcriptome of *A. aerophoba* and *D. avara*, respectively.

**Table 1.** Biological replicates per condition and time point.

| Species | Treatment | Time | Replicates |
| --- | --- | --- | --- |
| *A. aerophoba* | Seawater microbial consortia | 1h | 4 |
|  |  | 3h | 5 |
|  |  | 5h | 5 |
|  | *A.aerophoba* symbiont consortia | 1h | 5 |
|  |  | 3h | 5 |
|  |  | 5h | 5 |
| *D. avara* | Seawater microbial consortia | 1h | 4 |
|  |  | 3h | 4 |
|  |  | 5h | 3 |
|  | *A.aerophoba* symbiont consortia | 1h | 4 |
|  |  | 3h | 3 |
|  |  | 5h | 3 |

### Transcriptomic response upon seawater microbial and symbiont consortia

Significant differentially expressed genes (DEGs) were defined by edgeR, using a threshold of log2|FC| ≥2 (i.e., 4-fold change) and FDR p-value < 0.005, as in previous studies (Pita et al., 2018; Wu et al., 2022). The DEGs were classified as up-regulated and down-regulated in the symbiont treatment when compared to the expression levels in the seawater microbial treatment. The results from the differential expression analysis in edgeR and the full Trinotate annotation reports for the DEGs can be found in Tables S3 to S6.

### *D. avara* differential response to microbial consortia involves immune- and ubiquitin-related genes

We detected a total of 28 DEGs between *D. avara* sponges exposed to seawater and *A. aerophoba*-symbiont consortia and most genes showed higher expression levels in the symbiont treatment (Fig. 1). The highest proportion of DEGs was detected at 5h (Fig. 1A). Blastp provided annotation for 39 % of the total 64 DEGs (Fig. 1B) and searched in the SigP database classified all of them as non-transmembrane signaling peptides (Table S4). Expression profiles of *D. avara* individuals treated with each type of microbial consortia were consistent and biological replicates clustered together at all time points (Fig. 1C).

**Fig. 1.**
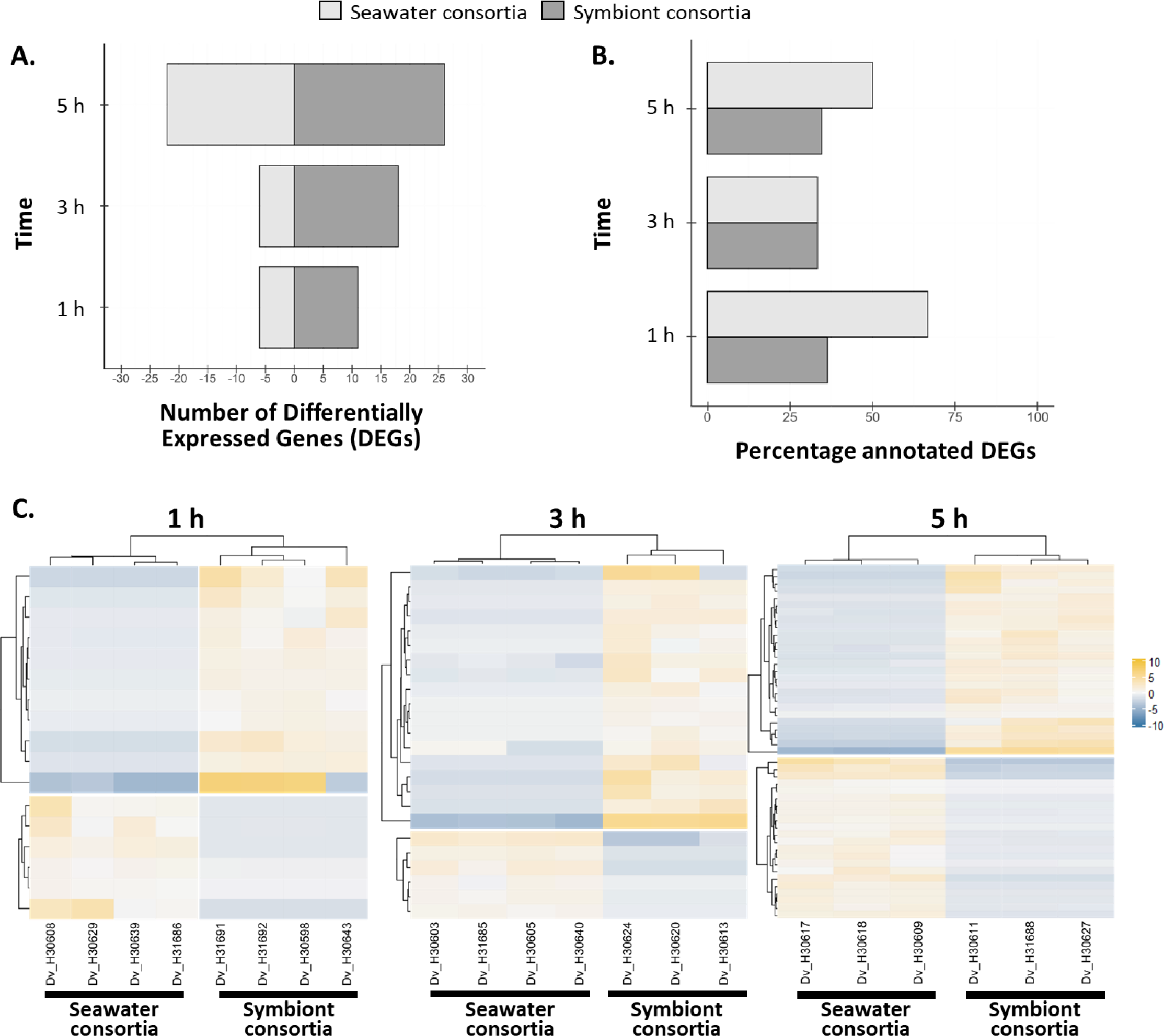
Differential gene expression of *D. avara* individuals treated with seawater microbial vs. *A. aerophoba*-symbiont consortia. **(A)** Number of differentially expressed genes (DEGs). Genes with increased expression upon symbiont encounter compared to seawater microbial consortia have positive values. **(B)** Percentage of DEGs with annotation for each microbial treatment and time point. **(C)** Heatmaps show the TMM-normalized relative expression level per DEG (rows) for each sample (columns) at 1 h, 3 h, and 5 h after microbial treatment. Genes were defined as differentially expressed with edgeR, FDR p-value < 0.005 and log2|FC|≥2.

Based on Pfam and blast annotations, we identified five differentially expressed genes encoding for NOD-like receptors (NLRs) (Fig. 2, within “immunity” category). Four out of five differentially expressed NLRs showed higher expression levels upon seawater microbes than upon *A. aerophoba*-symbiont exposure. Three of these (*TRINITY_DN18609_c0_g1, TRINITY_DN65570_c0_g1, TRINITY_DN18609_c0_g2*) were expressed at all time points and corresponded to incomplete NLRs (only the LRR-domain was detected; PF13516), and annotated as NLRC3 based on Blastp, whereas the fourth gene (*TRINITY_DN6063_c1_g1*), found only at 5 h (Fig. 2), contained the characteristic NACHT domain of NLRs (PF05729) and a peptidase domain (PF00656), and was assigned to the NLRC4 family based on Blastp annotation (Table S4). In contrast, there was one NLR that showed elevated gene expression in sponges incubated with symbionts in all time points (*TRINITY_DN42758_c1_g2*); it contained a NACHT domain and was assigned to the NLRP3 family based on Blastp annotation (Fig. 2; and Table S4).

**Fig. 2.**
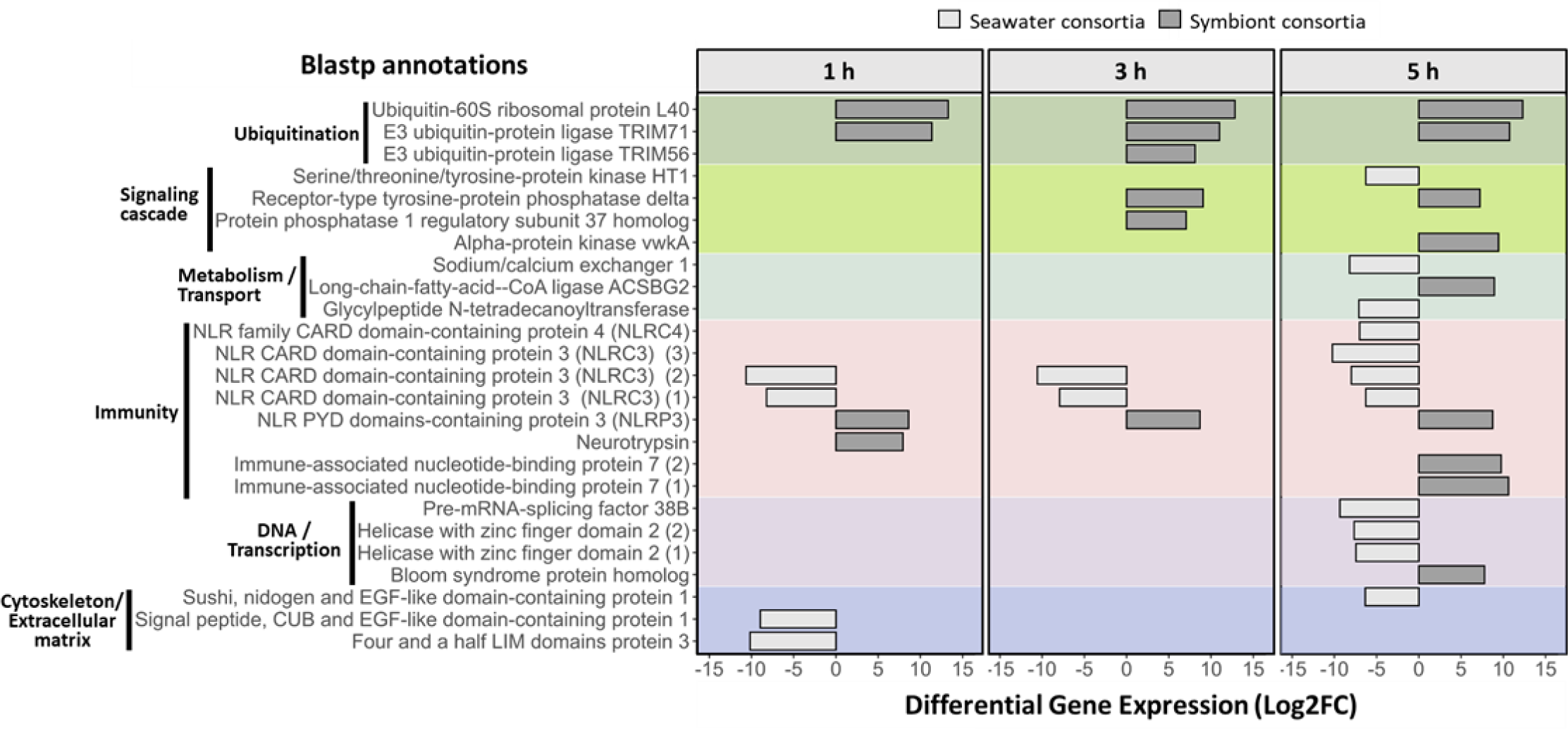
Functions and expression levels of differentially expressed genes in *D. avara* at 1 h, 3 h, and 5 h after seawater microbial and *A. aerophoba*-symbiont consortia treatment. Genes with increased expression upon symbiont encounter compared to seawater microbial consortia have positive Log2FC values. Genes were defined as differentially expressed with edgeR, FDR p-value < 0.005 and log2|FC|≥2. Only genes with Blast annotations are included. Numbers in brackets indicate different genes with the same annotation.

In addition to NLRs, we detected other DEGs potentially involved in innate immunity and ubiquitination that showed higher expression levels upon encounter to *A. aerophoba* symbionts than to seawater microbes (Fig. 2). Among the immune genes, we detected a SRCR-containing gene associated to neurotrypsin (*TRINITY_DN137847_c3_g1*; PF00530), and two genes related to an immune-associated GTP-binding protein (PF04548) (*TRINITY_DN1745_c0_g1* and *TRINITY_DN5077_c0_g1*) (Fig. 2; and Table S4). The regulation of ubiquitination was evident by the differential expression of three genes: one ubiquitin-60S ribosomal protein L40-like (*TRINITY_DN322946_c0_g1*) and two E3 ubiquitin ligases (*TRINITY_DN739_c0_g4* and *TRINITY_DN37530_c0_g1*, Fig. 2; and Table S4). The *A. aerophoba*-symbiont consortium also stimulated genes annotated as protein phosphatases (*TRINITY_DN4437_c1_g1* and *TRINITY_DN33881_c0_g1*) with fibronectin (PF00041) or LRR (PF13516) domains, an alpha-protein kinase (*TRINITY_DN61539_c1_g1*), a CoA ligase (*TRINITY_DN2791_c1_g1*), and a DEAD box-containing protein (*TRINITY_DN9624_c0_g1*; PF00270) (Fig. 2; and Table S4).

The seawater microbial treatment showed higher expression levels of genes involved in cell surface and cytoskeleton organization than compared to the symbiont consortia treatment, including a calmodulin-ubiquitin and epidermal growth factor-like containing gene (*TRINITY_DN5241_c0_g1*), and a LIM domain-containing gene (*TRINITY_DN182729_c0_g1*) (Fig. 2 and Table S4). At 5h, genes related to functions such as DNA regulation and transcription, metabolism and transport, and signaling cascades showed elevated gene expression levels in seawater microbial treatment. For example, two genes (*TRINITY_DN9504_c0_g1* and *TRINITY_DN41910_c0_g1*) for helicases with a zinc finger domain (HELZ2) belonging to the superfamily of P-loop NTPases (PF13087 and PF04851) which are predicted to be nuclear co-activators of the peroxisome proliferator-activated receptors (Fig. 2 and Table S4) were identified. We also detected two genes (*TRINITY_DN9504_c0_g1* and *TRINITY_DN41910_c0_g1*) involved in the molecular function of calcium and calmodulin binding. One of these genes contained a nidogen-like domain (PF06119), which is predicted to enable Notch binding activity and to be involved in cell-matrix adhesion, whereas the other gene with a Calx-beta motif (PF03160) regulates the transport of calcium and sodium across the cell membrane. In addition, a serine/threonine tyrosine-protein kinase (*TRINITY_DN63_c1_g1*), a glycylpeptide N-tetradecanoyltransferase (*TRINITY_DN16852_c2_g2*) involve in lipid modification, and an mRNA-splicing factor (*TRINITY_DN150336_c1_g1*) showed also higher gene expression levels 5 h after seawater consortia treatment than in symbiont treatment (Fig. 2 and Table S4).

### *A. aerophoba* differential response to microbial consortia involves signaling genes, ubiquitination-related genes and kinases

Differential gene expression between *A. aerophoba* sponges incubated with seawater microbial and symbiont consortia was only observed at 5 h, (log2|FC| ≥2 (i.e., 4-fold change) and FDR p-value < 0.005), (Fig. 4A) and showed consistently elevated expression profiles in seawater microbial treatment than symbiont treatment (Fig. 4B). Within the total 11 DEGs detected, 9 genes were signaling peptides, as reported by signalP (and two contained a transmembrane domain Table S6). We identified three genes with additional blast annotation. A leucine-rich repeat receptor like protein kinase (*TRINITY_DN146410_c5_g2*) with similarity to a *Dictyostelium discoideum* gene (YTYK2; DDB_G0283397), an ubiquitin ligase (*TRINITY_DN163315_c3_g2*; LIN41), and a transposase-derived protein antagonist of heterochromatin (*TRINITY_DN169091_c1_g1*; ALP1) (Fig. 3B). If relaxing the significance threshold (log2|FC| ≥1 (2-fold change) and FDR p-value < 0.05), the number of DEGs increased but the pattern of elevated expression levels of DEGs in the seawater microbial consortia than symbiont treatment was consistent (Fig. S2).

**Fig. 3.**
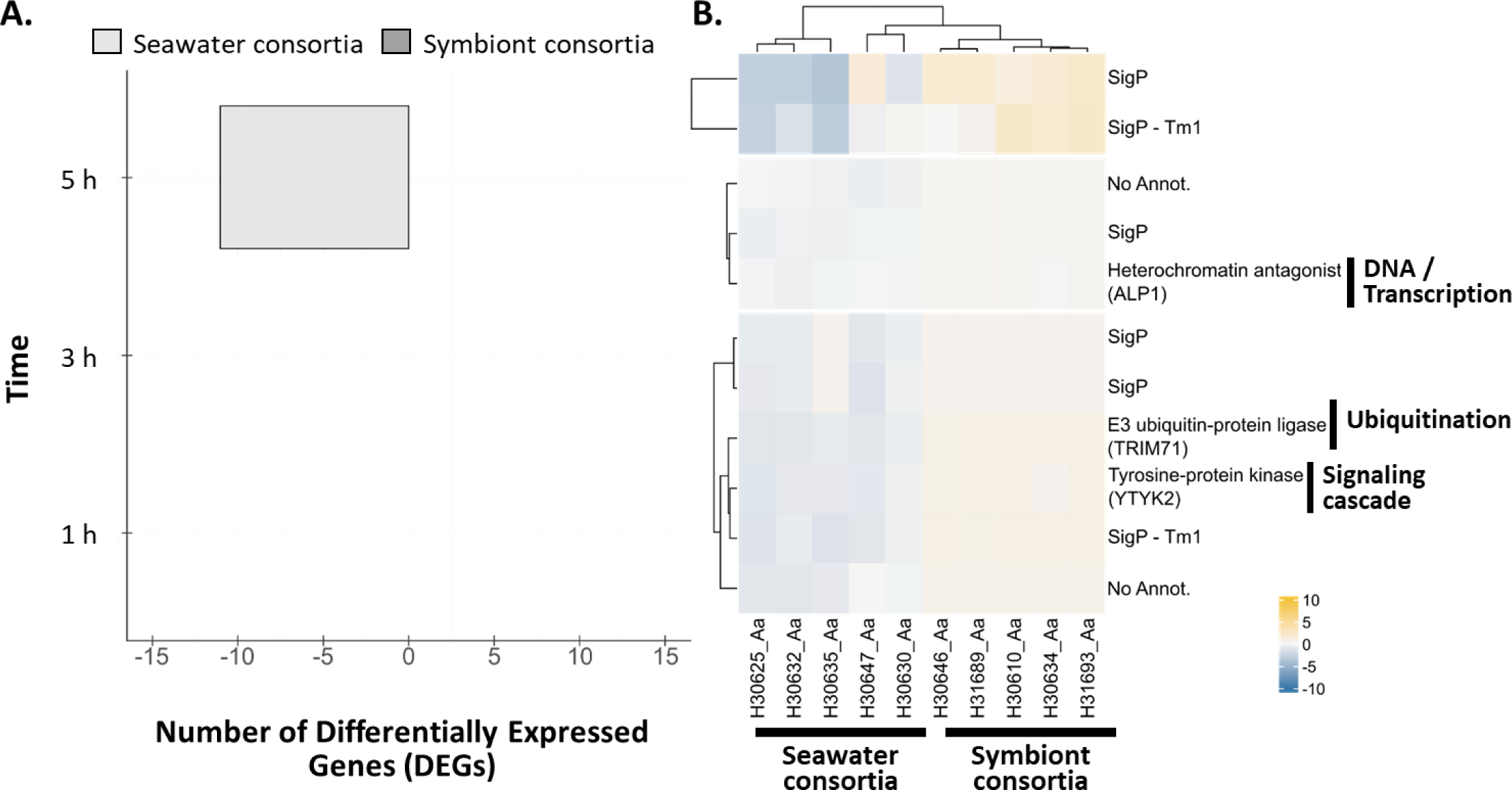
Differential gene expression of *A. aerophoba* individuals treated with seawater microbial consortia vs. sponge own symbionts. **(A)** Number of differentially expressed genes (DEGs). Genes with increased expression upon symbiont encounter compared to seawater microbial consortia have positive values. **(B)** Heatmap show the TMM-normalized relative expression level per DEG (rows) for each sample (columns) at 5 h after microbial treatment. Functions of DEGs are included only for genes with Blast annotations (right bold legend). Genes were defined as differentially expressed with edgeR, FDR p-value < 0.005 and log2|FC|≥2.

**Fig. 4.**
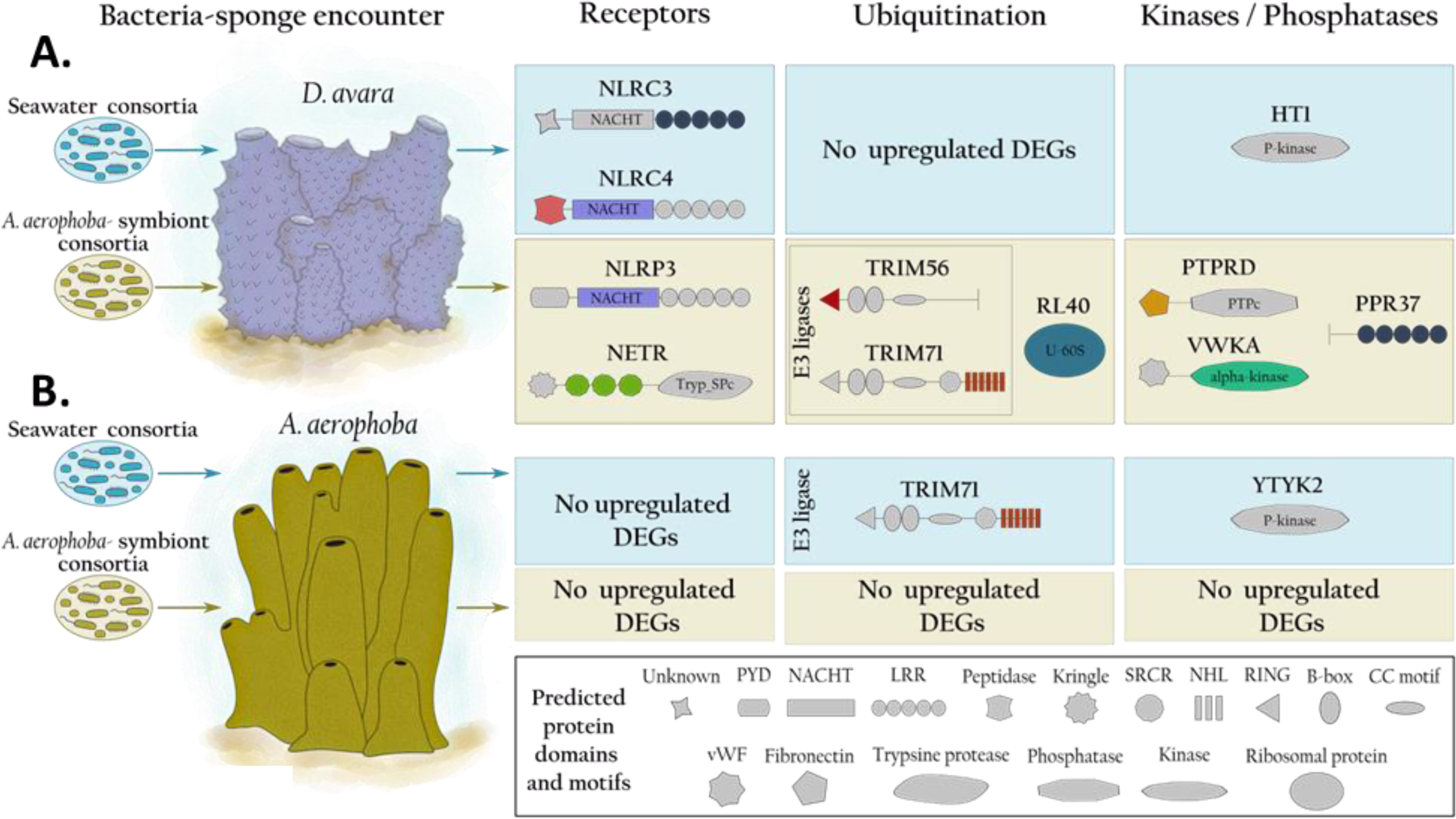
Overview of the transcriptomic response in **(A)** *D. avara* and **(B)** *A. aerophoba* upon microbial consortia encounter, derived either from seawater or symbiont preparations. DEGs detected after each treatment and annotated as receptors or related to ubiquitination and signaling are shown along with UniProt ID of best blastp hits. Colored domains had a Pfam annotation, whereas gray-shaded ones were not detected, and the potential structure of each gene was drawn based on smart.embl.de.

## Discussion

In this study, we characterized the transcriptomic responses of the Mediterranean sponges *A. aerophoba* and *D. avara* upon incubation with either microbial seawater or *A. aerophoba*-symbiont consortia. Previous studies comparing filtration rates showed that sponges take up seawater bacteria at higher rates than symbiotic bacteria (Wehrl et al., 2007; Wilkinson et al., 1984). Among transcriptomic studies in which sponges were subject to different stimuli (Koutsouveli et al., 2020; Pita et al., 2018; Posadas et al., 2021; Schmittmann et al., 2021; Wu et al., 2022), this one is among the first to simulate natural conditions and consequently, the overall host responses involved moderately fewer DEGs. The sponge species investigated responded differently to the experiment. The LMA sponge *D. avara* discriminated between treatments via NLR receptors, whereas no PRRs were differentially regulated in the HMA sponge *A. aerophoba*. This is the first time that the differential regulation of various NLR families is linked to microbial discrimination in sponges. While differentially-expressed genes in *D. avara* showed higher levels of expression upon symbiont consortia encounter than to seawater microbial consortia, little differential expression between both treatments was observed in *A. aerophoba*. We propose that the way sponges distinguish between microbes may depend on the HMA-LMA status as well as on species-specific traits.

### Moderate transcriptional response of sponges to microbial exposure

The exposure of *A. aerophoba* and *D. avara* to seawater microbial consortia and *A. aerophoba-* symbionts showed differential gene expression of few genes (i.e., < 70 genes) for both sponge species investigated (Fig. 1A and Fig. 3A), even when significance threshold was relaxed (Fig. S2). In the current study, we detected a relatively lower transcriptional response (i.e., number of DEGs) than in previous studies, particularly for *A. aerophoba.* A previous experiment assessing the response of both sponge species studied here to commercial microbial elicitors (i.e., lipopolysaccharide and peptidoglycan), compared to a sham injection with filtered artificial seawater, detected > 400 DEGs and ca. 49 DEGs in *A. aerophoba* and *D. avara*, respectively (Pita et al., 2018). In another study, the transcriptional response of *A. aerophoba* to wounding included thousands of DEGs (Wu et al., 2022). Furthermore, *A. queenslandica* juveniles in response against its native bacteria compared to foreign bacteria involved the differential gene expression of >1000 genes (Yuen, 2016). Besides potential differences due to bioinformatics analysis, we propose that, to some extent, the different magnitude of differential gene expression observed in previous, and this study are linked to the microbial stimuli applied (commercial vs. “natural”), the way they were presented to the sponge (injection vs. incubation), and the life stage (juveniles vs. adults) of the sponges. It thus appears that the magnitude of the host response is scalable depending upon the treatment, ranging from low (natural bacterial consortia), to medium (commercial elicitors), to high (mechanical damage). Figuratively speaking, in this study we are listening to the sponge “whispering”, as opposed to “talking” upon injection with elicitors (Pita et al., 2018), and even to “screaming” upon mechanical damage and snail predation (Wu et al., 2022). Furthermore, the higher number of immune related genes between in *A. queenslandica* juveniles compared to adult *D. avara* and *A. aerophoba* individuals could also be related to the maturation of the sponge immune system. The development and acquisition of immunity remains to be studied in sponges, but in other organisms (e.g., zebrafish, honey bees, mice and humans) a series of maturation steps are required for achieving immunocompetence, and this competence is adapted to the different life stages and previous encounters of the host with microbial cues (Gätschenberger et al., 2013; Lam et al., 2004; Park et al., 2020; Yang et al., 2015).

The present study presented the microbes in a way that is closer to natural conditions, in which sponge filter-feeding lifestyle translates into constant encounter with microbes in the surrounding water. The maintenance and implementation of immune mechanisms is energetically demanding, and the physiological costs may represent trade-offs between other metabolic activities (Ardia et al., 2012; Palmer, 2018). Thus, we speculate that the immune activity is constitutive in adult sponges due to their constant interactions with microbes. In fact, constitutive expression of a great variety of PRRs has been observed in the sponge *Halichondria panicea* (Schmittmann et al., 2021). Induced responses will be activated upon other type of stimuli, like “damage signals”, as in the response of *A. aerophoba* to wounding (Wu et al., 2022). In contrast, the response to microbes is based on “fine-tuning” of immune components that are already in place. The constant interactions of sponges with their microbiome and seawater bacteria including potential pathogens may favor constitutive expression over induced activation of immune components. This strategy challenges traditional views on induced immunity from terrestrial animals, but it may indeed be widespread among marine invertebrates (Schmittmann et al., 2021; Williams & Gilmore, 2022).

### Sponges recognize microbial consortia differently

*D. avara* individuals incubated with *A. aerophoba*-symbiont consortia showed an overall approx. 50 % higher number of DEGs compared to *A. aerophoba* sponges (Fig. 1A and 3A). We observed a differential expression of immune receptors in *D. avara* such as NLRs (Fig 2 and Fig. 4A), whereas no PRRs were differentially expressed in *A. aerophoba* (Fig. 3B and Fig. 4B). Moreover, *A. aerophoba* showed a lower transcriptomic response to symbiont (“self”) than to seawater microbial consortia (“non-self”). The few genes differentially regulated in *A. aerophoba*, showing mainly reduced expression levels upon its own symbiont treatment (Fig. 3B), could indicate that the sponge detects microbes in a different way than *D. avara*. The low transcriptomic response in *A. aerophoba* could be the result of the sponge recognizing its own symbionts as “self” or of its HMA status. Since we did not detect high differential gene expression between the microbial treatments in *A. aerophoba*, we argue this to be related to the sponge HMA status rather than with “self”/“non-self” recognition. Ideally, we would have included a fourth treatment consisting of *D. avara*-symbiont consortia exposure to clarify these hypotheses, but we were not able to extract symbionts from *D. avara* in sufficient quantity for our incubations, due to its LMA status. Moreover, we cannot exclude the possibility that *A. aerophoba* sponge cells were remaining in the microbial symbiont fraction and, thus, affected the transcriptional responses. However, we would expect that those sponge cells will activate a transcriptional response in both *D. avara* and *A. aerophoba*, if any, because studies on sponge self- and non-self-transplants suggest active rejection of cells from other sponges, even if derived from other individuals of the same species (Hildemann et al., 1979; Hirose et al., 2021; Saito, 2013). Overall, our results show a lack of a differential transcriptomic response in a HMA sponge against microbes that we interpret as an adaptation to the permanent presence of symbionts within the sponge mesohyl system. More sponge species representative of the HMA-LMA categories will however be needed to validate and elaborate this hypothesis.

### *D. avara* and *A. aerophoba* employ different sets of genes

The differential transcriptomic response of *D. avara* individuals to symbionts and seawater microbial consortia involved several immune genes including one SRCR-containing receptor and two GTP-binding proteins. The genes encoding for the immune-associated GTP-binding protein (IAN) in *D. avara* are part of the GIMAP family. IAN genes were not detected in any other early divergent metazoans including the genome of the sponge *A. queenslandica*, but are broadly and patchy distributed among eukaryotes, and orthologs have been reported in plants, corals, and molluscs as means of microbial defense (Coelho et al., 2022; Liu et al., 2008; McDowell et al., 2016; Weiss et al., 2013). GIMAP family is conserved among vertebrates, where it is implicated in the development and maintenance of immune cells (e.g., lymphocytes (Limoges et al., 2021)). SRCRs are involved in the recognition of a broad range of ligands and are highly diversified in invertebrates (Buckley & Rast, 2015; Neubauer et al., 2016; Smith et al. 2018). These receptors are also expanded in sponges (Pita et al., 2018; Ryu et al., 2016; Schmittmann et al., 2021) and potentially play a role in sponge symbiosis. For instance, a SRCR-domain containing gene was up-regulated in symbiotic individuals of the sponge *Petrosia ficiformis* compared to aposymbiotic individuals (i.e., photosymbiont-free), suggesting the involvement of this immune receptor in sponge symbiosis (Steindler et al., 2007). Moreover, in juveniles of *A. queenslandica,* different SRCRs were upregulated in response to native and foreign bacteria (Yuen, 2016). Altogether, SRCRs arise as putative mediators of sponge-microbe interactions in different sponge species and the GIMAP family may as well deserve more attention in future studies.

In comparison to *D. avara*, the lower differential transcriptomic response of *A. aerophoba* was limited to reduced expression of a kinase-like receptor, an E3 ligase and an antagonist of heterochromatin in symbiont vs. seawater microbial consortia treatment (Fig 3B and Fig. 4B). The antagonist of like-heterochromatin protein (ALP1) in plants acts as a transposase mediating various cellular pathways and capable of silencing gene expression involving E3 ubiquitin ligases (Golbabapour et al., 2013; Ohtsubo et al., 2008). The role of this transposase in inhibiting transcriptional responses is proposed to have evolved as a means for evading surveillance by the hosts (Liang et al., 2015). In fact, pathogens cause a variety of transcriptional changes (e.g., alteration of chromatin structure, proteolytic degradation, deactivation of transcription factors, etc.) to exploit a wide range of pathways which enhances their survival within the host (De Monerri & Kim, 2014; Villares et al., 2020). We therefore hypothesize that the late (i.e., at 5h) down-regulation of both ALP1 and E3 ubiquitin ligase in symbiont compared to seawater microbial exposure (Fig. 3B) could indicate active host gene silencing by symbionts, to prevent becoming target material for degradation. Functional studies are imperative to validate the processes in which the detected DEGs are involved, yet this remains a challenge in sponges as models for genetic manipulation are currently limited to explants or cells of two sponge species (Hesp et al., 2020; Revilla-I-Domingo et al., 2018).

### *D. avara* distinguishs between seawater microbial and symbiont consortia via NLRs

*D. avara* sponges differentiated between seawater microbes and symbionts via differential expression of NLRs. Moreover, the differentially-expressed NLRs with higher expression levels in sponges incubated with seawater microbial consortia were similar to the NLRC3 and NLRC4 families (based on Blastp results; Fig 2 and Fig. 4A), whereas the NLR that exhibited higher expression levels in response to *A. aerophoba* symbionts showed similarity to the NLRP3 family (Fig 2 and Fig. 4A). A phylogenetic analysis of these NLRs was not possible because our transcripts for these NLR-like genes were incomplete (i.e., lacking NACHT or LRR domains, Fig 4A). To confirm if these NLRs belonged to different subfamilies, we performed an additional blast search (at protein level, e-value < 1e−5; Table S7) of the differentially expressed NLRs in *D. avara* against the freshwater sponge *Ephydatia muelleri*, for which a chromosome-level genome is available (Kenny et al., 2020). The best hits of differentially-expressed *D. avara* NLRs in *E. muelleri* support that the NLRs activated in response to seawater bacteria belong to a different NLR subfamily than the one responding to *A. aerophoba* symbiont consortia (Table S7).

Differential gene expression of NLRs in *D. avara* was accompanied by differential expression of ubiquitination, kinases and phosphatases (Fig 2 and Fig. 4A). We speculate that the different types of NLRs (i.e., NLRC3, NLRC4 and NLRP3) activate different downstream signaling pathways in *D. avara* whose ultimate goal is to regulate microbial recognition by the sponge. In humans and mice these NLR families regulate inflammatory pathways (Pan et al., 2022; Schneider et al., 2013; Sun et al., 2022; Uchimura et al., 2018; Walle & Lamkanfi, 2016). Inflammation requires various post-translational modifications comprising ubiquitin ligases, kinases and phosphatases (Akther et al., 2021; Song & Li, 2018; Yang et al., 2017), and the ubiquitin system, which is crucial in many biological process, is proposed as an essential innate immunity regulator and as a modulator of host-microbe interactions (Li et al., 2016; Zhang et al., 2021). Overall, our results show experimentally for the first time the role of NLRs in microbial discrimination by sponges and suggest their role in sponge-symbiont interactions. Importantly, LMA sponges are known to contain an expanded and diverse set of NLRs compared to HMAs (e.g., (Germer et al., 2017; Ryu et al., 2016; Schmittmann et al., 2021; Yuen et al., 2014)). The regulation of NLRs in the LMA sponge *D. avara*, compared to the non-regulation of these receptors in the HMA sponge *A. aerophoba*, may further support the previous hypothesis that the HMA-LMA status may influence how the sponge immune system responds to microbial encounters. The first experimental evidence of enhanced NLRs expression in sponges was reported in *D. avara* as a response to commercial microbial elicitors (Pita et al., 2018). Our results build on these observations and support the participation of poriferan NLRs in specific microbial recognition. Future studies should focus on identifying the ligand of this different NLRs to finally provide functional evidence of the role of sponge NLRs in immune specificity.

### Conclusion

The molecular mechanisms employ in early divergent metazoans for microbial discrimination are still only poorly understood. In the present study, we characterized the transcriptomic response of two sponge species upon incubations with seawater microbes and sponge-derived symbionts by RNA-Seq differential gene expression analysis. Our observations showed that sponges mount a moderately low (less than70 DEGs) but different transcriptomic response to natural microbial encounters. Microbial discrimination in sponges seems to be driven by the repertoire of immune genes harbored by the host and the degree in which these are induced. The HMA sponge *A. aerophoba* showed little differential gene expression and no participation of PRRs upon microbial exposure. Contrastingly, our results support the involvement of NLRs in specific microbial discrimination in the studied LMA sponge. We hypothesize that the different NLR families under regulation might trigger various signaling pathways in *D. avara* which are tuned to recognize among different microbial cues. Furthermore, we suggest that the differential response to microbial exposure between sponge species could be the result of species-specific traits or HMA-LMA features that influence the regulation of immune components in the host. To unveil more potential sponge molecular adaptations to microbial encounters it is crucial to investigate different sponge species along the LMA-HMA spectrum, under experimental setups that resemble as much as possible natural conditions, and to test different microbial structures that may induce or silence the transcriptomic response of the host. Finally, conducting comparative studies between the relevant genes mediating microbial discrimination in sponges and other early divergent invertebrates would further expand our understanding on the role of PRRs on microbial recognition and place sponges with their unique life-styles in an evolutionary context.

## Supplementary Material

Supplementary data are available at Genome Biology and Evolution online.

## Acknowledgments

We are grateful to Rafel Coma and Manel Bolívar (CEAB-CSIC) for assistance during the sponge collection. We thank Marc Catllà and the personal from the ZAE at ICM-CSIC for assistance during the experimental work in Barcelona. We acknowledge the staff from IKMB sequencing facilities for cDNA library preparation and sequencing. We also thank Dr. Lara Schmittmann and Dr. Vasiliki Koutsouveli for helpful discussions and feedback in the data analysis. LP received supported by “la Caixa” Foundation (ID 10010434), co-financed by the European Union’s Horizon 2020 research and innovation program under the Marie Sklodowska-Curie grant agreement No 847648), fellowship code is 104855. LP and MR received additional institutional support by the “Severo-Ochoa Centre of Excellence” accreditation (CEX2019-000928-S). This is a contribution from the Marine Biogeochemistry and Global Change research group (Grant 2021SGR00430, Generalitat de Catalunya). UH was supported by the DFG (“Origin and Function of Metaorganisms”, CRC1182-TP B01) and the Gordon and Betty Moore Foundation (“Symbiosis in Aquatic Systems Initiative”, GBMF9352).

## Data availability

The raw reads, metadata, transcriptome assembly and full annotation for this study have been deposited in the European Nucleotide Archive (ENA) at EMBL-EBI under the accession number PRJEB61959 (ERP147040).

## Supplementary information

**Text S1.** Characterization of microbial consortium treatments by flow cytometry

The concentration of the microbial consortia stocks obtained by enrichment was estimated via flow cytometry and adjusted to reach 10^5-6^ bacteria mL^-1^ final concentration in each experimental aquarium. In addition, water samples (2 mL) from each aquaria were collected right before the experiment (T-1h) and right after (T0h) adding the microbial consortium. Samples for flow cytometry were fixed in paraformaldehyde + glutaraldehyde (1% + 0.05% final, respectively) and stored at -80°C until analysis. Microbial cell concentration was quantified by flow cytometry (FACSCalibur, Becton-Dickinson, 488 nm excitation blue laser) following the method of Gasol and Morán (1999). In short, DNA in microbial cells was stained with Syto13, and detected based on cell-side scatter, forward scatter, and green fluorescence of the stained DNA. Plastic beads were used as reference for plotting. Bacterial cell concentrations were calculated based on number of events and calibrated flow rate.

Although the aquaria were kept overnight in 1 µm-filtered seawater and an additional 0.1 µm-filter was applied for 3 h before the experiments, the water in the aquaria was not sterile, some bacterial cells remained (Fig. S1 A and C). We could still detect the addition of the treatment, particularly in the cell population of higher DNA content, in both seawater and *A. aerophoba* symbiont treatments (Fig. S1 B and D, R6 gate). We could detect and increment of one order of magnitude in the bacterial concentrations in the water before and after the addition of microbial consortia, to a final concentration ∼10^6^ cells/mL.

**Fig. S1.**
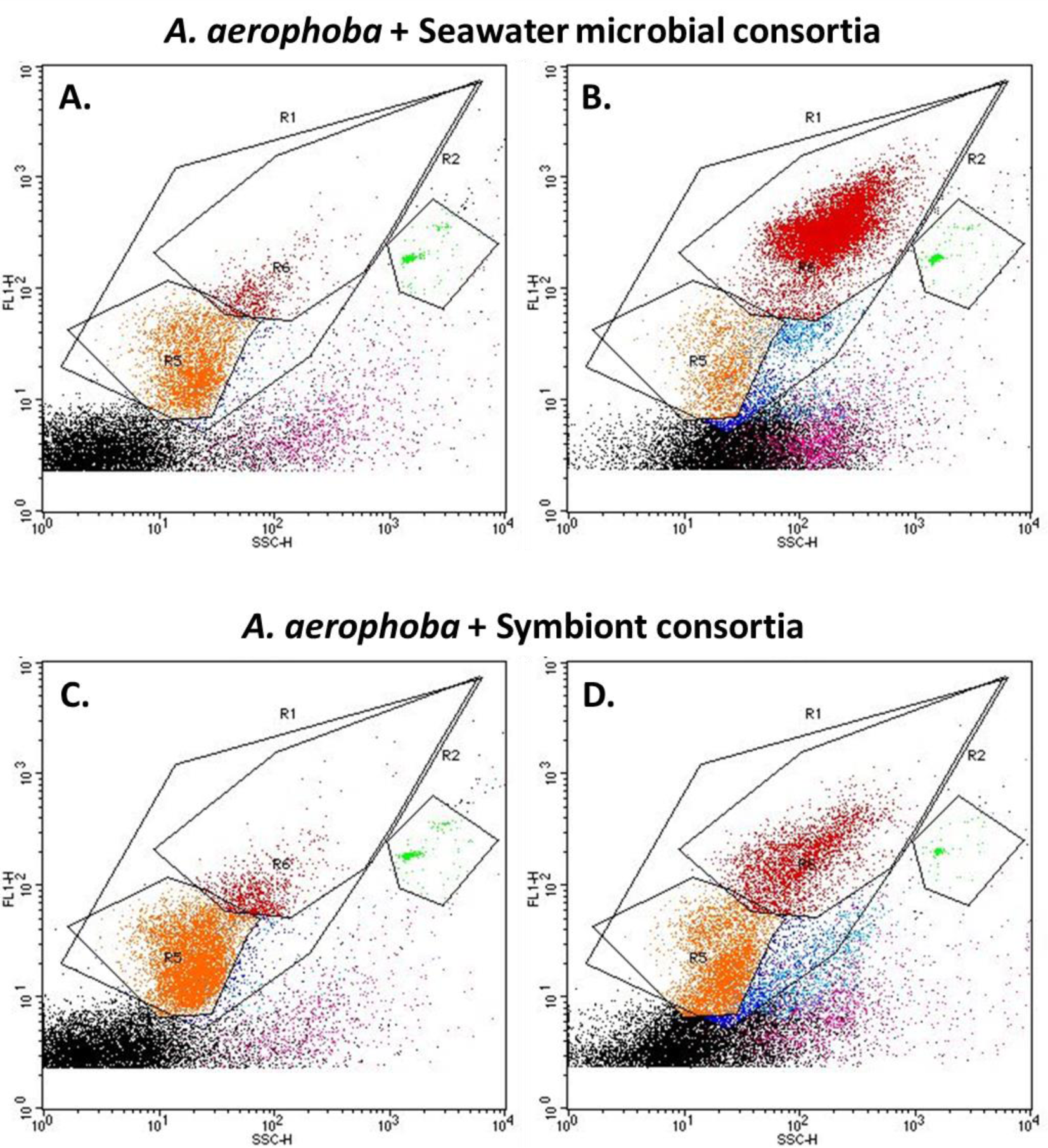
Representative cytograms of seawater consortia **(A-B)** and *A. aerophoba*-symbiont **(C-D)** consortium used for the experiments. The microbial stock concentration was estimated before (T-1) **(A-C)** and after (T0) **(B-D)** adding the bacteria to the incubation tank. R1: all bacteria; R5 and R6: low and high DNA bacteria, respectively; R2: quantification beads. Water samples (2 mL) from all aquaria were collected before the experiment (time point -1h) and every hour during the course of the experiments (time points 0, 1, 2, 3, 4, 5 h), and fixed in paraformaldehyde + glutaraldehyde (1% + 0.05% final, respectively). Microbial cell concentration in the water by was quantified by flow cytometry (FACSCalibur, Becton-Dickinson, 488 nm excitation blue laser) following the method of Gasol and Morán (1999), to assess the sponge filtration activity. The bacterial cells were stained with Syto13, and detected based on cell-side scatter, forward scatter, and green fluorescence of the stained DNA. For comparison with the sponges, control aquaria (i.e., without sponge) were also exposed to the microbial treatments and sampled at the same time points.

**Table S1.** Number of read pairs (million reads). “Raw” refers to the output from sequencing; “Clean” to surviving read pairs after trimming and filtering in trimmomatic-v0.38; and “Eukaryote” to pairs identified as non-prokaryotic and nonmicrobial eukaryote by kaiju-v1.6.2 (Menzel & Krogh, 2015).

| Average per library<br>(± standard error) | Raw | Clean | Eukaryote |
| --- | --- | --- | --- |
| *A. aerophoba* | 23.8 ± 1.8 | 22.1 ± 9.2 | 15.0 ± 6.3 |
| *D. avara* | 19.6 ± 1.2 | 18.1 ± 10.7 | 11.7 ± 0.7 |

**Table S2.**
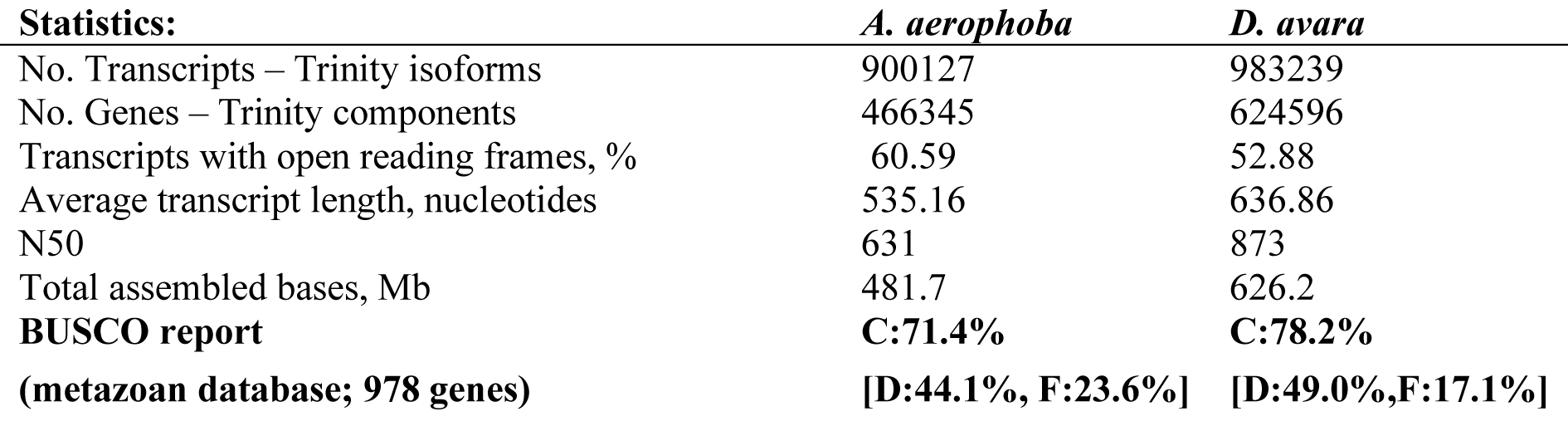
Statistics of the *de novo* transcriptomic assemblies. Transcripts refer to Trinity isoforms, genes refer to Trinity components. Mb: mega bases.

**Table S3.** Differential Expression analysis for *D. avara* at as identified in edgeR (FDR p-value < 0.005 and log2|FC|≥2) at 1h, 3h and 5h (Excel file)

**Table S4.** Annotation of the differentially expressed genes for D. avara identified in edgeR (FDR p-value < 0.005 and log2|FC|≥2) at 1h, 3h and 5h (Excel file)

**Table S5.** Differential Expression analysis for A. aerophoba at as identified in edgeR (FDR p-value < 0.005 and log2|FC|≥2) at 5h (Excel file)

**Table S6.** Annotation of differentially expressed genes for A. aerophoba identified in edgeR (FDR p-value < 0.005 and log2|FC|≥2) at 5h (Excel file)

**Table S7.** Blastp results of *D. avara* differentially expressed NLRs against *Ephydatia muelleri* (e-value < 1e−5) (Excel file)

**Fig. S2.**
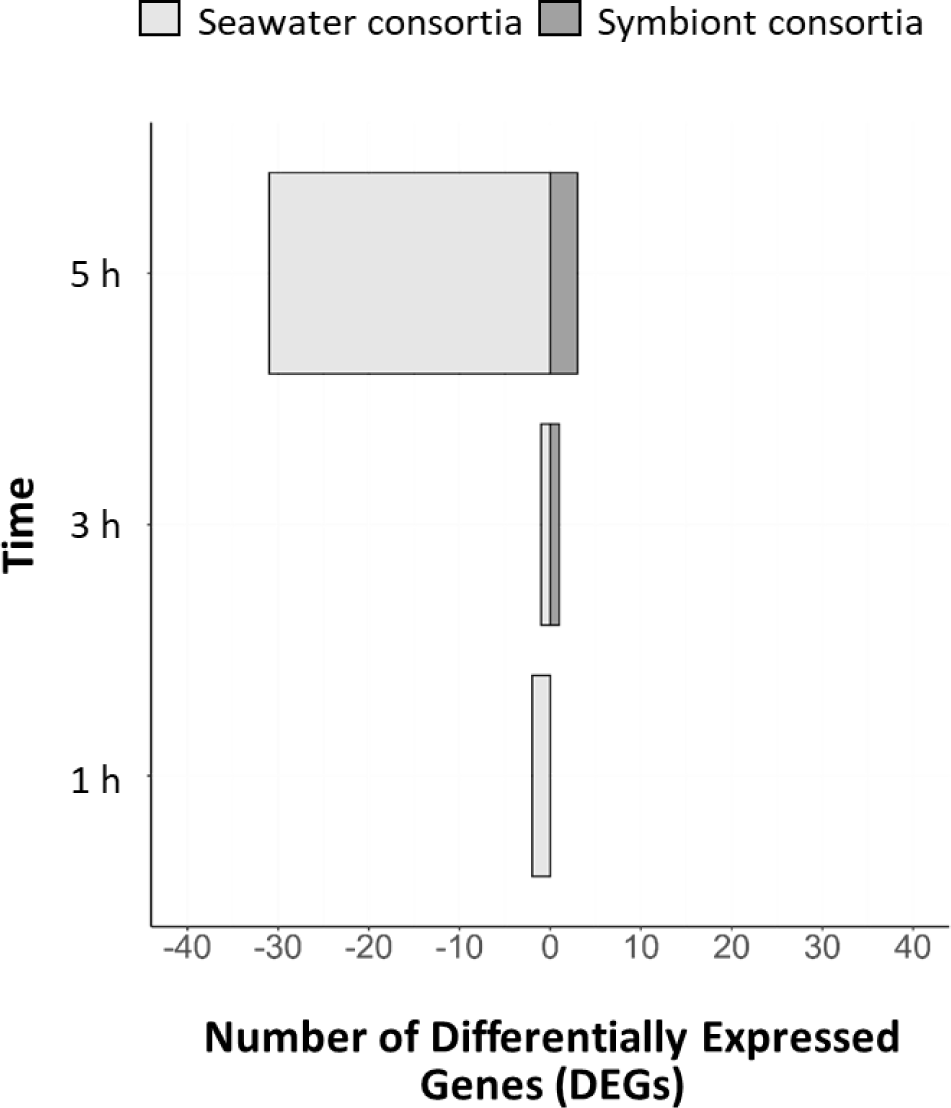
Number of differentially expressed genes (DEGs) of *A. aerophoba* individuals treated with seawater microbial consortia vs. symbiont consortia. Negative values reflect number of genes with lower expression levels in seawater microbial consortia treatment compared to symbiont treatment. Genes were defined as differentially expressed with edgeR, FDR p-value < 0.05 and log2|FC|≥1.

